# Gray matter volume increase in the retrosplenial/posterior cingulate cortices of blind soccer players

**DOI:** 10.1101/2024.09.04.611145

**Authors:** Tomoyo Morita, Eiichi Naito

## Abstract

Spatial navigation is a crucial brain function that occurs when an individual moves from one location to another. This function normally relies on vision, and the retrosplenial/posterior cingulate cortices (RSC/PCC) and parahippocampal cortex (PHC) play crucial roles. However, an extraordinary adaptation can be observed in blind soccer players, where the human brain can perform spatial navigation without vision. Therefore, this study tested the possible increase in gray matter (GM) volume in the RSC/PCC and PHC in the brains of a world’s top blind soccer player and other blind soccer players. We collected structural magnetic resonance imaging (MRI) scans from six blind soccer players (including the world’s top player) and eight blind non-soccer players. Using voxel-based morphometry (VBM) analysis (single-case VBM approach), we examined GM volume increase in each participant compared to 250 sighted participants (all of whom had never played blind soccer). Compared with the sighted participants, the world’s top blind soccer player had a significant increase in GM volume in the bilateral RSC/PCC. Two of the other five blind soccer players also showed an increase in the left RSC/PCC. However, such an increase in GM volume was not observed in the blind non-soccer players. Eventually, the probability of a significant increase in the RSC/PCC was significantly higher in the blind soccer group than in the blind non-soccer group. In contrast, only one blind soccer player (not the top player) showed a significant increase in the PHC, and no between-group difference was observed in the probability of a significant increase in the PHC. This study, unveiling the characteristics of the brains of the world’s top blind soccer player and of other blind soccer players, for the first time demonstrates that blind soccer training, which requires spatial navigation based on non-visual cues, may enlarge the human RSC/PCC and provide findings that promote understanding the brains of challenged persons playing blind soccer.

## 1 Introduction

Spatial navigation is a crucial brain function when a person moves from one location to another and normally relies on vision. However, the human brain sometimes exhibits extraordinary adaptations to this function in the absence of vision. A typical example can be observed in blind soccer players. A world’s top blind soccer player exists who can move to the right place where he wants to be on the soccer court as if he can see (Blind Football World Cup YouTube 2018). To realize this remarkable performance, this player’s brain must use multisensory (e.g., auditory, vestibular, motor, and proprioceptive) information other than vision and have developed the ability for spatial navigation without vision.

In sighted individuals, several meta-analyses of spatial navigation tasks have consistently reported activations in the medial parietal region immediately behind the splenium, caudal part of the corpus callosum, and parahippocampal cortex (PHC; Epstein, 2008; Spreng et al., 2009; Boccia et al., 2014; Cona and Scarpazza, 2019; Li et al., 2021; Baumann and Mattingley, 2021). The medial parietal region anatomically corresponds to the ventral posterior cingulate cortex (PCC); however, it is conventionally referred to as the retrosplenial cortex (RSC; Foster et al., 2023). Therefore, in this study, we refer to this section as RSC/PCC. Since damage to the RSC/PCC or PHC region usually causes wayfinding deficits (Takahashi et al., 1997; Suzuki et al., 1998; Aguirre and D’Esposito, 1999; Barrash et al., 2000), they are believed to play a crucial role in spatial navigation in sighted people.

Regarding spatial navigation without vision, the inactivation of the RSC in rats trained in a maze task in light has been shown to cause behavioral impairments in the task in darkness rather than in normal light (Cooper and Mizumori, 1999; Cooper et al., 2001). This suggests the particular importance of the rat RSC in spatial memory and navigation, especially in the absence of visual cues. However, it is not known whether human PCC/RSC also have the same function. On the other hand, little is known about whether the PHC is involved in spatial navigation without vision in both humans and animals. If the human RSC/PCC and PHC also play crucial roles in spatial navigation based on non-visual cues, these regions in the top-blind soccer player must play essential roles for his remarkable ability in spatial navigation.

Currently, it is well established that in experts (e.g., athletes and musicians) with specific training over a long period, gray matter (GM) volume increases in their brain regions that repeatedly process information related to the content of the training (Maguire et al., 2000; Gaser and Schlaug, 2003; Draganski et al., 2004; Kanai and Rees, 2011; Thomas and Baker, 2013; Gerber et al., 2014). If the world’s top-blind soccer player relies on the RSC/PCC and PHC for spatial navigation without vision, we may expect a GM volume increase in these brain regions.

Another important question is whether the expected GM volume increase in these regions may also occur in other blind soccer players or even in visually impaired individuals with no blind soccer experience (blind non-soccer). By addressing these questions, we can discuss the generalization of the possible GM volume increase in the RSC/PCC and PHC among blind soccer players and the importance of blind soccer experience for GM volume change.

Therefore, we collected structural magnetic resonance imaging (MRI) scans from six blind soccer players (including the world’s top player), eight blind non-soccer players, and 250 sighted individuals (with no blind soccer experience). Using voxel-based morphometry (VBM) analysis (single-case VBM approach), we examined GM volume increase in the RSC/PCC and PHC of the six blind soccer and eight blind non-soccer players compared to the 250 sighted individuals.

## 2 Materials and Methods

### 2.1 Participants

Fourteen visually impaired males (age range: 22–42 years) participated in this study. They included six persons who had played blind soccer (BS group; mean age, 31.3 ± 6.0 years) and eight with no blind soccer experience (BNS group; mean age, 33.3 ± 7.7 years). One participant (BS1) in the BS group was the world’s top blind soccer player, aged 26 years. The participant lost his sight at the age of 8 years and has been playing blind soccer since the age of 10 years. He was a member of the Brazilian blind soccer national team and has competed in four consecutive Paralympic Games, contributing to Brazil’s gold medal victories in each event. The other four members of the BS group were current or past members of the Japanese blind soccer national team (BS2, BS3, BS4, and BS5). However, the remaining participant was not a member of the national team but had many years of blind soccer experience (BS6). Details of the blind participants, including their clinical features, are summarized in Table 1.

**Table 1.**
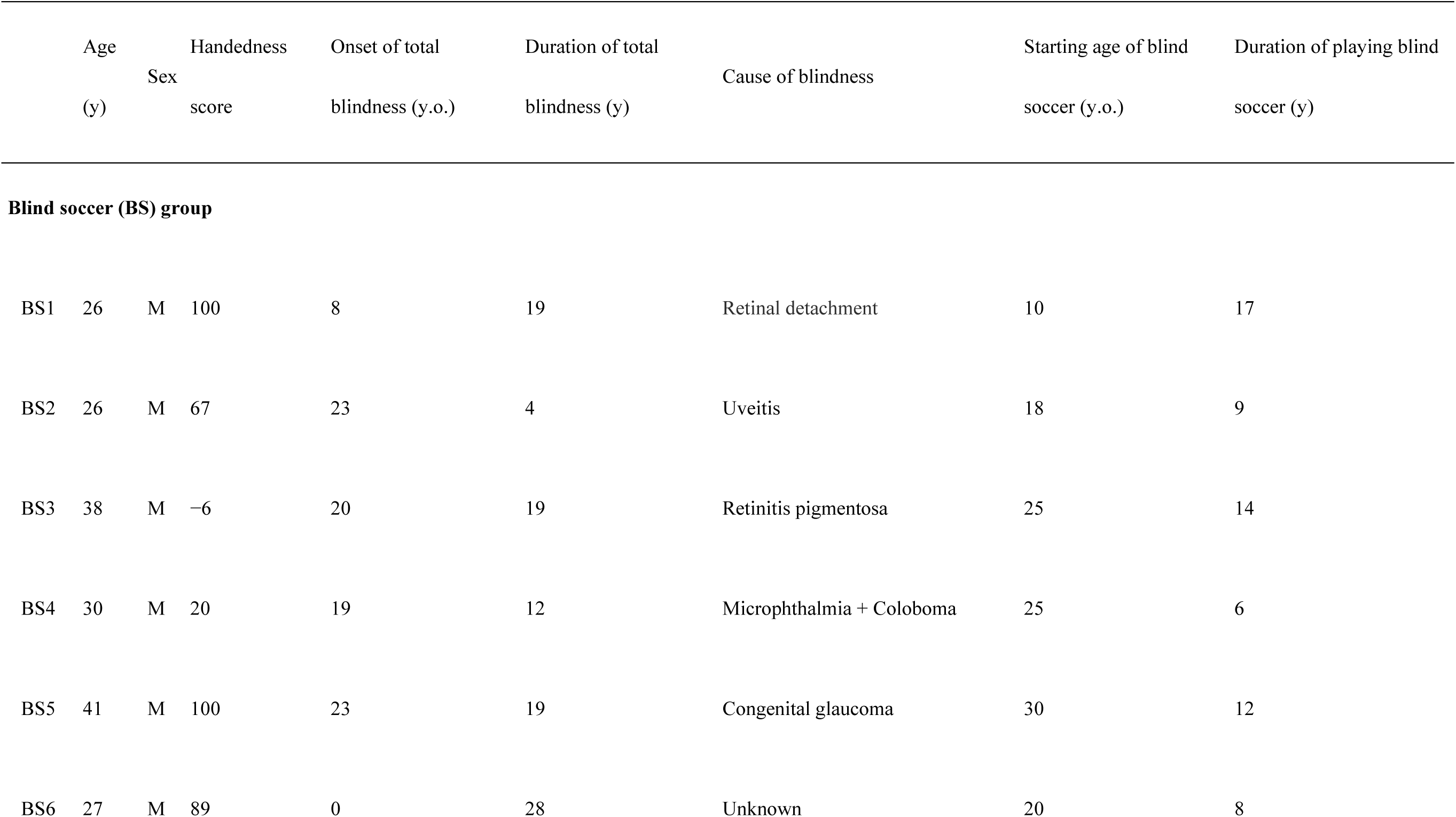
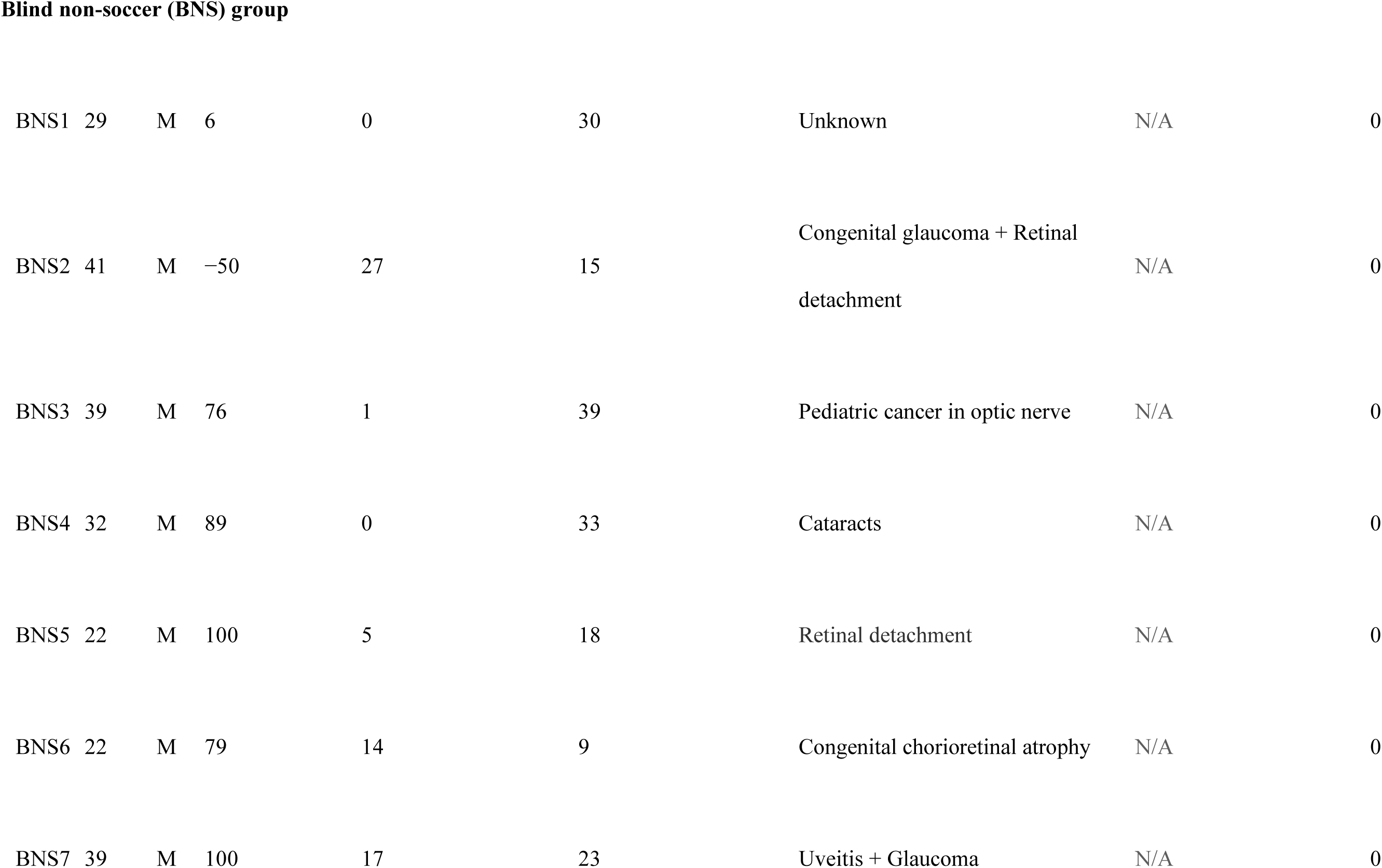
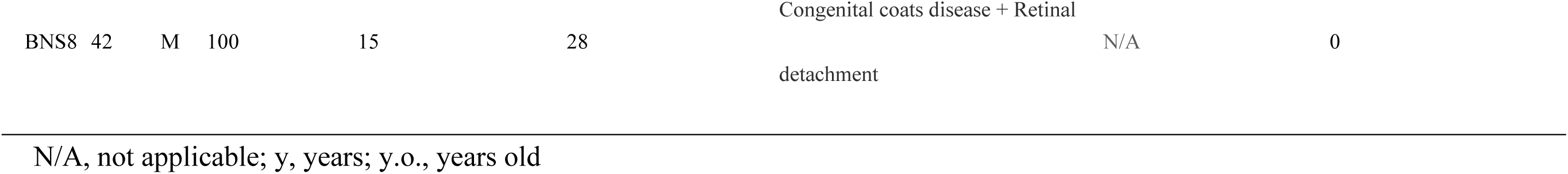
Visually impaired participants: summary of their clinical features and blind soccer experience.

In total, 250 sighted male participants with no experience in blind soccer were also involved in this study (sighted group; mean age, 24.1 ± 5.4 years, range: (21–54 years). The age range of the sighted participants was set to include that of the visually impaired participants, although the mean age of the sighted participants was slightly lower than that of the visually impaired participants. 236 of 250 sighted participants were asked how many years they had been doing what kind of sports and their sports training experience for more than 1 year continuously in the past. 76 of 236 participants reported they had more than 1 year experience of soccer, and 53 participants reported they had more than 4 years experience. None of the participants had any history of psychiatric disorders. Handedness was confirmed using the Edinburgh Handedness Inventory (Oldfield, 1971). The mean handedness scores were 61.6 (range: -6 to 100) in the BS group, 62.6 (range: -50 to 100) in the BNS group, and 71.1 (range: -100 to 100) in the sighted group.

The study protocol was approved by the Ethics Committee of the National Institute of Information and Communications Technology and the MRI Safety Committee of the Center for Information and Neural Networks (CiNet; no. 2003260010). The details of the study were explained to the participants before the start of the experiment., and they provided written informed consent. In cases where the visually impaired participants could not provide their signatures, we obtained their oral informed consent and written informed consent signed by an experimenter or their guardians in their presence. This study was conducted in accordance with the principles of the Declaration of Helsinki (1975).

### 2.2 MRI acquisition

For each participant, a T1-weighted magnetization-prepared rapid gradient echo (MP-RAGE) image was acquired using a 3.0-Tesla MRI scanner (Trio Tim; SIEMENS, Germany) and a 32-channel array coil. Imaging parameters were as follows: repetition time, 1,900 ms; echo time, 2.48 ms; flip angle, 9°; field of view, 256 × 256 mm^2^; matrix size, 256 × 256 pixels; slice thickness, 1.0 mm; voxel size, 1 × 1 × 1 mm^3^; and contiguous transverse slices, 208.

### 2.3 Preprocessing of MRI data

VBM analysis was performed to examine the change in GM volume in each visually impaired participant compared with that in sighted participants (single-case VBM). First, we visually inspected the T1-weighted structural images of all participants and confirmed the absence of observable structural abnormalities and motion artifacts. Data were analyzed using Statistical Parametric Mapping (SPM 12; Wellcome Centre for Human Neuroimaging, London, UK) running on MATLAB R2018b (MathWorks, Sherborn, MA, USA). The following analytical procedures were performed as recommended by Ashburner (Ashburner, 2010) and used in our previous study (Morita et al., 2022).

The T1-weighted structural image of each participant was segmented into GM, white matter (WM), cerebrospinal fluid (CSF), and non-brain parts based on a tissue probability map provided by the SPM. Through this procedure, segmented GM and WM images were generated. These images were approximated to the tissue probability map using rigid-body transformation and transformed GM and WM images were generated.

Diffeomorphic Anatomical Registration Through Exponentiated Lie Algebra (DARTEL) processing was performed to generate a DARTEL template that was subsequently used to accurately align the segmented images across participants. In this process, the average of the transformed GM and WM images across all participants was defined as a template that included the GM and WM. A deformation field was calculated to deform the template into the transformed image for each participant, and the inverse of the deformation field was reapplied to the individual image. This series of operations is performed multiple times to generate a sharply defined DARTEL template.

Thereafter, an affine transformation was applied to the DARTEL template to align it with the tissue probability map in the Montreal Neurological Institute (MNI) standard space. The segmented GM image of each participant was then warped nonlinearly to the DARTEL template represented in the MNI space (spatial normalization). Next, the warped image was modulated by the Jacobian determinants of the deformation field to preserve the relative GM volume, even after spatial normalization.

Finally, the modulated GM image of each participant was smoothed with an 8-mm full-width-at-half-maximum Gaussian kernel and resampled to a resolution of 1.5 × 1.5 × 1.5 mm voxel size. The smoothed image was used for statistical inference.

### 2.4 Statistical analysis

To compare the GM volume between each visually impaired participant and the 250 sighted individuals, a two-sample non-parametric permutation test (without variance smoothing) was performed using the Statistical non-Parametric Mapping toolbox (version 13.1.09; Nichols and Holmes, 2002). Non-parametric statistics provide a valid alternative to parametric statistics in the context of single-case VBM, as they do not rely on the assumptions of normal distribution or equal variance (Scarpazza et al., 2016). For each statistical comparison, we adopted 251 permutations based on the sighted group size of 250. Age, handedness scores, and total brain volume (i.e., the sum of GM and WM volumes) were entered into the design matrix as covariates of no interest to minimize the impact of these variables on the results.

Since we had structural hypotheses for the RSC/PCC and PHC, we conducted a region-of-interest (ROI) analysis. We prepared an ROI for each of the left and right RSC/PCC and PHC based on a previous study that conducted a meta-analysis of functional neuroimaging studies on spatial navigation (Cona and Scarpazza, 2019). Regarding the RSC/PCC, we initially created a sphere of 12 mm radius around the peak in the left or right RSC/PCC reported in this study (left RSC/PCC peak coordinates x, y, and z = -12, -50, and 18; right RSC/PCC peak coordinates x, y, and z = 14, -45, and 18). We selected this radius to ensure that the spheres are located anterior to the parieto-occipital sulcus without extending into the occipital cortex. However, these spheres partially overlapped the corpus callosum, which is a WM region. Therefore, we created a GM mask image from the current data using the SPM Masking Toolbox (Ridgway et al., 2009). By applying this image, we exclusively depicted truly positive GM voxels within each sphere and defined left or right RSC/PCC ROI.

A similar step was performed for the PHC. A meta-analysis study (Cona and Scarpazza, 2019) reported a peak only in the right PHC (peak coordinates x, y, and z = 24, -40, and -10). Therefore, we created a sphere of 12 mm radius around this peak. For the left PHC, we created a sphere of 12 mm radius around the right-to-left flipped peak (peak coordinates x, y, and z = -24, -40, and -10) of the right peak and defined putative left PHC. However, these spheres partially overlapped with regions anatomically defined as other than the PHC (e.g., the hippocampus). Therefore, the “parahippocampal” region of the Automatic Anatomical Labelling Atlas (AAL; Tzourio-Mazoyer et al., 2002) was used as the anatomical map of the PHC. Eventually, the PHC ROI in each hemisphere was defined by depicting the overlap between the sphere and anatomical map in each hemisphere.

Within each ROI, we examined the increase in GM volume in each visually impaired participant compared to that in the sighted group. We searched for significant clusters of voxels (cluster-forming threshold, p < 0.005) within each ROI (extent threshold, p < 0.05, Family-Wise Error-corrected). We also counted the number of participants in the BS and BNS groups who showed significant clusters within each ROI and evaluated the between-group differences in this proportion using Fisher’s exact test. In each group, we summed the total number of participants with significant clusters in either the left or right ROI for the left and right RSC/PCC and PHC ROIs.

To visualize the individual GM volumes in the identified clusters, we calculated the GM volume of each cluster in each participant (Fig. 1). The GM volume was divided by the individual total brain volume to adjust for the head size of each participant. These values were plotted strictly for visualization purposes, and no statistical evaluation was performed to avoid the circular assessment issues raised by Kriegeskorte et al. (2009).

**Figure 1.**
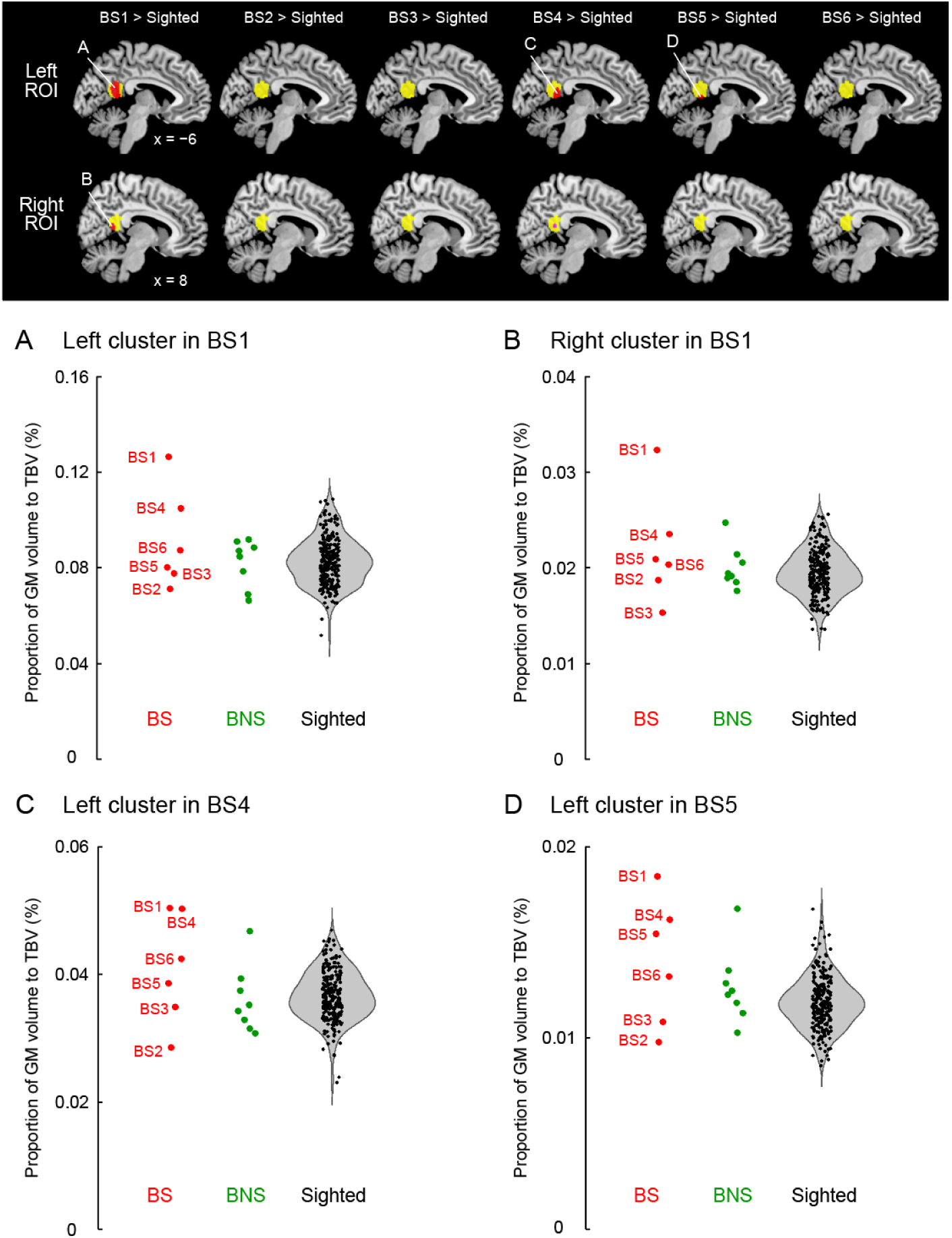
Results from single-case VBM analysis. The upper panel shows significant clusters of voxels indicating GM increase (red sections) within the left and right RSC/PCC ROIs (yellow sections) compared to the 250 sighted participants. BS1 showed a significant cluster in each ROI (red section, A: left, B: right). B4 and B5 showed a significant cluster in the left ROI (red sections, C, D), and B4 showed a cluster with significant trend in the right ROI (pink section, p = 0.08). These sections are superimposed on the MNI standard brain, and the sagittal slices (x = -6 and 8) are shown. (A-D) Graphs show individual GM volumes in each of the identified RSC/PCC clusters. A vertical axis indicates the proportion of GM volume in the cluster to TBV. Red, green, or black dots represent the data obtained from the participants of the BS, BNS, or sighted groups, respectively. These plots are horizontally jittered to avoid over-plotting. Violin plots are also shown for the data obtained from the sighted group. Abbreviations: BS, blind soccer; BNS, blind non-soccer; GM, gray matter; MNI, Montreal Neurological Institute; PCC, posterior cingulate cortex; ROI, region of interest; RSC, retrosplenial cortex; TBV, total brain volume; VBM, voxel-based morphometry.

## 3 Results

### 3.1 RSC/PCC

Compared to the 250 sighted participants, BS1 showed significant clusters of voxels where the GM volume increased both in the left (peak coordinates x, y, and z = -6, -51, and 14; T = 4.14; 551 voxels) and right (peak coordinates x, y, and z = 14, -45, and 9; T = 3.78; 184 voxels) RSC/PCC ROIs (Fig. 1A, B). These were parts of one large bilateral RSC/PCC cluster (1034 voxels), which was the largest cluster in the whole brain. A similar increase in GM volume was observed in the other two blind soccer players. BS4 and BS5 had significant clusters of voxels in the left RSC/PCC ROI (BS4: peak coordinates x, y, and z = -12, -44, and 8; T = 4.56; 244 voxels; Fig. 1C, BS5: peak coordinates x, y, and z = -3, -51, and 9; T = 4.20; 82 voxels; Fig. 1D). In addition, BS4 showed another cluster with significance trend in the right RSC/PCC ROI (peak coordinates x, y, and z = 5, -48, and 15; T = 2.85; 43 voxels; p = 0.08). In contrast, none of the blind non-soccer players showed substantial clusters (> 10 voxels) within the RSC/PCC ROIs. Eventually, the probability of significant clusters within the left and right RSC/PCC ROIs (4/12) in the BS group was significantly higher than that (0/16) in the BNS group (Supplementary Table 1, Fisher’s exact test, p = 0.02).

When we checked the individual GM volumes in each identified cluster, BS1 showed the largest value among all participants, which deviated from the distribution of sighted participants (Fig. 1A-D).

### 3.2 PHC

For the PHC, BS1 did not show significant clusters within the left or right PHC ROI. Only BS4 showed a significant cluster in the left PHC ROI (peak coordinates x, y, and z = -20, -30, and -8; T = 3.84; 72 voxels; not shown in the figure). In the BNS group, none of the participants showed significant clusters, and BNS3 merely showed a cluster with significant trend in the right PHC ROI (peak coordinates x, y, and z = 27, -39, and -3; T = 3.79; 46 voxels). Eventually, the probability of significant clusters within the left and right PHC ROIs (1/12) in the BS group was not significantly different from that (0/16) in the BNS group (Supplementary Table 2; Fisher’s exact test, p = 0.43).

## 4 Discussion

### 4.1 General

In this study, a direct evaluation of the spatial navigation ability without the vision of blind soccer players was missing. However, the hyperspatial navigational ability of BS1 is evident (Blind Football World Cup, YouTube, 2018), and the function of spatial navigation has likely been trained through blind soccer training in other blind soccer players. The GM volume increased in the bilateral RSC/PCC in the world’s top blind soccer player (BS1) and the left RSC/PCC in the other two blind soccer players, although such a significant increase was not observed in the blind non-soccer players. Eventually, the BS group had a significantly higher probability of a significant increase in RSC/PCC than the BNS group. In contrast, GM increase in the PHC was rare, and no between-group difference was observed. Furthermore, no GM increase in the RSC/PCC is reported in sighted individuals with regular soccer training (Li et al., 2024), which is also confirmed in the present sighted participants with soccer experience (see Supplementary Materials and Supplementary Fig. 1), rebuffing the view that the present GM increase in the RSC/PCC is owing to general soccer training. Viewed collectively, the series of results suggest that blind soccer training, which requires spatial navigation based on non-visual cues, may increase the human RSC/PCC volume and imply that the RSC/PCC is more important than the PHC during blind soccer.

The current work was a single-case VBM study of challenged individuals with blindness. We adopted this approach rather than a group analysis because (1) it allowed us to depict distinct features of GM volume at the individual level when compared to the control group, and (2) the mean of a small sample (for example, Six individuals for the BS group and eight for the BNS group) is usually affected by outliers largely. However, the small number of visually impaired participants is a limitation of this study. This makes it difficult to conclude what percentage of GM volume in a significantly larger population of blind soccer players is increasing their RSC/PCC and whether such an increase is indeed absent in a larger population of blind non-soccer players. However, our study provides novel evidence that some blind soccer players, represented by BS1, showed an extraordinary increase in GM volume in the RSC/PCC, which should be referred to as hyper-adaptation, as we have proposed in studies of a top wheelchair racing Paralympian (Morita et al., 2022; 2023). Such hyperadaptation is presumed to be achieved through highly motivated, long-term daily training while overcoming the difficulties of disability (Supplementary Fig. 2).

The exact physiological changes underlying the increase in GM volume remain unknown. Axon sprouting, dendritic branching synaptogenesis, neurogenesis, and changes in the glial number and morphology have been suggested as important contributors (Zatorre et al., 2012). An increase in GM volume in one brain region is usually use-dependent and is related to the duration (amount) of a particular training (Maguire et al., 2000) and associated skill level (Gaser and Schlaug, 2003; Gerber et al., 2014; James et al., 2014). The greatest GM volume increase in the RSC/PCC (Fig. 1) of BS1 appears to fit these views because his training period (17 years) was the longest (Table 1), and his spatial navigation ability was outstanding. However, since no significant increase was observed in BS3, who had the second-longest training period (14 years), and a significant increase was observed in BS4, who had the shortest training period (6 years), the training period was not the only determining factor. The onset of total blindness was at the age of 8 years for BS1, while those for the Japanese-blind soccer national team members (BS2, BS3, BS4, and BS5) were approximately at the age of 20 years. Therefore, only BS1 started blind soccer training since puberty (Table 1). Long-term training from puberty may be advantageous for morphological changes (Morita et al., 2022; 2023; Steele et al., 2013). Finally, the congenital blind soccer player (BS6) showed no significant increase; however, it remains unclear whether congenital or acquired blindness could be another factor affecting GM changes. Therefore, the extension of this study to a relatively larger population of blind soccer players will provide important insights into the factors that may affect plastic GM changes in the RSC/PCC.

### 4.2 Roles of RSC/PCC in blind soccer players

The RSC/PCC is composed of the following two major subregions: granular (Brodmann area [BA]29) and dysgranular (BA30) areas (Vann et al., 2009). Most of the RSC/PCC sections where BS1 showed an increase in GM volume appeared to overlap with BA30, which is a posterior subregion of the RSC/PCC. A study combining diffusion tensor imaging and functional connectivity (Li et al., 2018) has shown that the superior parietal lobe (SPL), PHC, and visual cortices, which are the structures involved in spatial navigation with vision (Baumann and Mattingley, 2021; Vann et al., 2009), are anatomically and functionally connected to BA30.

A higher ability for spatial navigation without vision is advantageous during blind soccer because this ability allows players to move from one location to another more precisely within the court. When sighted individuals play futsal (indoor or salon soccer), they can immediately recognize and update their spatial locations within the same-sized court by simply looking at their surrounding environment. However, this is difficult for blind soccer players because they cannot use visual information. To recognize and update their spatial locations within the court, they must rely on auditory cues generated from their surrounding environment and the voices of callers and coaches instead of vision. In addition, they rely more on path integration, in which the brain integrates self-movements over time to update its current location by utilizing vestibular, motor, and proprioceptive information. Finally, this egocentric (self-centered) sensory-motor information must remain associated with allocentric (world-centered) information about the spatial map of the court to update their spatial locations within the court.

Among the abovementioned cortical regions involved in spatial navigation with vision (Vann et al., 2009), egocentric information has been proposed to be processed in the parietal cortex, whereas allocentric information is processed in the PHC and visual cortices. The RSC/PCC is situated in a region that connects these areas and, therefore, would be an appropriate place for the association between egocentric and allocentric information (Vann et al., 2009; Clark et al., 2018). In addition, the RSC/PCC is believed to be involved in this association function (Byrne et al., 2007; Iaria et al., 2007; Epstein, 2008). A patient with a hemorrhage in the right RSC may have difficulty in performing a task that requires the transformation and integration of egocentric and allocentric information (Suzuki et al., 1998).

This assumption appears to be supported by the functional connectivity results obtained from our additional functional MRI experiments. In the series of the current work, we also measured brain activity using the same scanner, while the same 14 visually impaired participants and 16 of the 250 sighted individuals performed an imaginary spatial navigation task (see details in Supplementary materials and Amemiya et al. (2021)). The task was a mental rehearsal of walking or jogging around a circle of 2 m radius, which was previously experienced without vision on a different day. Among the visually impaired participants, only BS1 showed clusters with significant trends (p = 0.059 corrected) in the early visual cortices (peak coordinates x, y, and z = -8, -84, and 16 in area hOc1[V1]; T = 5.37; 65 voxels) and right SPL (peak coordinates x, y, and z = 14, -52, and 74 in area 5 L; T = 7.17; 115 voxels), in which activity enhanced functional coupling with that in the left RSC/PCC ROI (seed) during the task, compared to the sighted participants (see Supplementary Fig. 3). It appears that when BS1 performs mental walking or jogging, the RSC/PCC is likely to communicate with the SPL (area 5) with an egocentric framework (Scheperjans et al., 2005; 2008; Naito et al., 2008) and with the visual cortices innately having an allocentric (retinotopic) framework (Striem-Amit et al., 2015), which corroborates the above view that the RSC/PCC is situated in an appropriate place for the association between egocentric and allocentric information. These facts allow us to hypothesize that BS1 repeatedly recruits the RSC/PCC for the association between egocentric and allocentric information when moving within a space without vision.

The enhanced functional coupling between the RSC/PCC and the visual cortices in BS1 is worth discussing. After the functional MRI experiment, BS1 reported that he could visualize the walking or jogging scene during the task, as well as the court scene in which he is playing blind soccer. Therefore, it would be worth while to study whether such functional coupling will be associated with the mental visualization of movement in space in the future.

### 4.3 Parahippocampal and hippocampal cortices

Previous neuroimaging studies on spatial navigation have consistently reported activation in the PHC in addition to the RSC (Epstein, 2008; Spreng et al., 2009; Boccia et al., 2014; Cona and Scarpazza, 2019; Li et al., 2021), suggesting that these are important brain regions for spatial navigation in sighted individuals. In our study, the results suggest that blind soccer training may significantly affect GM volume changes in the RSC/PCC but not in the PHC. This indicates that the RSC/PCC and PHC have different roles, as suggested by previous studies (Epstein, 2008; Epstein and Baker, 2019), and that spatial cognitive functions trained through blind soccer training do not require the PHC. The PHC is involved in scene information processing in sighted individuals and represents allocentric information about visual scenes (Vann et al., 2009). Visually impaired individuals cannot process such bottom-up visual scene information. However, unlike the visual cortices, where GM atrophy is observed in visually impaired individuals (Noppeney et al., 2005: Pan et al., 2007; Ptito et al., 2008; Leporé et al., 2010; Jiang et al., 2015), no GM atrophy has been reported in the PHC. Therefore, visually impaired participants might utilize the PHC for information processing other than bottom-up visual scene processing, which is not specifically required for playing blind soccer. Further studies are required to elucidate the role of PHC in these individuals.

In contrast to the PHC, visually impaired individuals (both congenital and acquired) are known to have an enlarged hippocampal volume (especially on the right side) compared to sighted individuals (Fortin et al., 2008; Leporé et al., 2009). Consistent with previous findings, the present visually impaired group (n = 14) showed a significant increase in GM volume in the right hippocampal ROI defined by the AAL, compared to the sighted group (Supplementary Materials and Supplementary Fig. 4). Such significant increase was not observed on the left side. We also performed a single-case VBM analysis for the left and right AAL-defined hippocampal ROI and found no significant differences between the BS and BNS groups (Supplementary Materials). Therefore, similar to the PHC, spatial cognitive functions trained through blind soccer training do not appear to require the hippocampus beyond its general roles in spatial memory and navigation in visually impaired individuals.

## 5. Conclusion

This study revealed that the volume of the RSC/PCC is significantly increased in the top blind soccer player and also in other blind soccer players compared to sighted individuals. This finding suggests that the RSC/PCC is involved in spatial navigation without vision, rather than the hippocampus and parahippocampus, believed to be important for spatial cognition in visually impaired individuals. This study has elucidated a role of human retrosplenial cortex by unveiling the neural underpinning for the outstanding spatial navigation ability of the world’s top blind soccer player and has greatly advanced our understanding the brains of challenged persons playing blind soccer.

## 7. Author Contributions

Both authors contributed to the conceptualization, validation, writing-review editing, investigation, funding acquisition, writing-original draft preparation, project administration, and supervision. TM: formal analysis and visualization. All authors have read and approved the final version of the manuscript, and agree with the order of presentation of the authors.

## Acknowledgments

The funding sources were not involved in the study design; collection, analysis, and interpretation of data; writing of the report; or the decision to submit the article for publication. The authors are grateful to Mr. Eigo Matsuzaki, Mr. Shigeo Murakami, and Mr. Kota Yamamoto of the Japanese Blind Football Association; Dr. Yasuo Nakamura of Doshisha University; Dr. Robert Nawa, Dr. Norikazu Sugimoto, Mr. Ryoma Yasuda, and Ms. Yoko Fujie of CiNet; Mr. Makoto Arimoto; Mr. Takanori Naruse of CR-NEXUS Inc.; and Mr. Masao Furuta of nac Image Technology for their invaluable support. The authors are also grateful to Dr. Satoshi Hirose, Dr. Tsuyoshi Ikegami, and Dr. Masaya Hirashima for their valuable help with the behavioral training.

## 9. Informed Consent Statement

Informed consent was obtained from all participants involved in the study. Informed consent has been obtained from the top blind soccer player (BS1) to publish this paper.

