## Supplementary materials for "Gray matter volume increase in the retrosplenial/posterior cingulate cortices of blind soccer players"

### No gray matter (GM) volume increase in the RSC/PCC of sighted individuals with soccer experience

236 of 250 sighted participants were asked how many years they had been doing what kind of sports and their sports training experience for more than 1 year continuously in the past. 53 of 236 participants reported they had more than 4 years experience, whereas 160 participants reported they did not have more than 1 year experience. We then examined brain regions that showed an increase in GM volume in the 53 participants, compared to the 160 participants. We performed a two-sample t-test for the whole brain. Age, handedness scores, and total brain volume (i.e., the sum of GM and WM volumes) were entered into the design matrix as covariates of no interest. When we used a voxel-wise threshold of $p < 0.005$ and a spatial extent threshold for a cluster of $p < 0.05$ (Family-Wise Error (FWE)-corrected), no significant clusters were identified. Therefore, we reported brain regions that showed GM volume increase in the entire brain (> 176 voxels [expected voxels per cluster, Friston et al., 1994]) in a voxel-cluster image using a height threshold of $p < 0.005$ uncorrected.

Most importantly, no voxels were identified in the bilateral RSC/PCC ROIs, which is compatible with a previous report (Li et al., 2024). This rebuffs the view that the present GM increase in the RSC/PCC is owing to general soccer training. Two clusters were identified in the entire brain (Red sections in Supplementary Fig. 1). One was located in the medial parieto-occipital cortices (area hOc6 [V6]: peak coordinates x, y, and z = -21, -71, and 23; T = 3.37; 442 voxels) extending into area hOc4d [V3A]. These dorsal occipital regions involve head-centered motion perception (Fischer et al., 2012; Schindler and Bartels, 2018) and spatial depth perception (Welchman, 2016; Ng et al., 2021). The other was in the cingulate motor area (peak coordinates x, y, and z = 2, -2, and 51; T = 3.25; 293 voxels).

### GM volume increase in the motivation-drive network of the visually impaired individuals

The wheelchair racing Paralympian study (Morita et al., 2022) reported an increase in white matter (WM) volume in the dorsomedial prefrontal region, a constituent of the motivation-drive network (Husain and Roiser, 2018). It was assumed that this phenomenon was related to the Paralympian's long-term daily training while overcoming the difficulties of the disability with higher motivation (Morita et al., 2022). This result led us to hypothesize that such a volume increase can also be observed in other individuals with disability, such as visually impaired persons. We then examined brain regions that showed an increase in GM or WM volume in the visually impaired group (n = 14) compared to the sighted group (n = 250). We performed a two-sample non-parametric permutation test for the whole brain. For this analysis, we used a voxel-wise threshold of $p < 0.005$ and determined significance in terms of the spatial extent of a cluster ($p < 0.05$, FWE-corrected).

The visually impaired group showed a significant increase in GM volume in the bilateral ventral striatum, including nucleus accumbens (NAc; peak coordinates x, y, and z = 0, 12, and -2; T = 4.65; 2354 voxels; red section in Supplementary Fig. 2). This is the largest and only significant cluster in the entire brain. The second largest cluster was found in the dorsomedial prefrontal cortex (dMPFC), extending into the dorsal anterior cingulate cortex (ACC) (peak coordinates x, y, and z = 6, 30, and 59; T = 4.95; 1582 voxels; pink section in Supplementary Fig. 2). This cluster had fairly large number of voxels; however, it did not reach significance after whole-brain correction. No significant increase in the WM volume was observed in any brain region.

The NAc is an important subcortical structure in the motivation-drive network (Husain and Roiser, 2018) that contributes to the transformation of motivation to action for rewards, working in concert with the medial frontal regions, such as the dMPFC and ACC (Gabbott et al., 2005; Hoover and Vertes, 2007; Husain and Roiser, 2018). Consistent with this view, lesions of the NAc profoundly affect motivation in animals, with a reduced willingness to allocate effort for rewards (Hauber and Sommer, 2009). Similarly, lesions and/or inactivation of the dMPFC impair the ability to work harder to receive greater rewards (Walton et al., 2003; Nakamura et al., 2022). In humans, significant bilateral NAc atrophy has been observed in apathetic patients with reduced motivation and passion (Carriere et al., 2014; Martinez-Horta et al., 2017). Based on these findings, one may assume that the increased GM volume in the NAc and dMPFC in the visually impaired group could be the outcome of their continuous challenges in taking daily actions with higher motivation despite the challenging circumstances of visual loss.

**Analysis of functional connectivity from the left RSC/PCC during an imaginary spatial navigation task**

BS1, BS4, and BS5 showed a significant increase in GM volume in the left retrosplenial/posterior cingulate cortices (RSC/PCC) (Fig. 1). Based on the previous finding that the superior parietal lobe (SPL), parahippocampal cortex (PHC), and visual cortices are anatomically and functionally connected to the RSC/PCC (Li et al., 2018), we examined whether the brain activity of these cortical regions enhanced its functional coupling with that of the RSC/PCC when a visually impaired participant performed an imaginary spatial navigation task compared to sighted participants by conducting a generalized psychophysiological interaction analysis (gPPI) implemented in the CONN toolbox (McLaren et al., 2012).

Procedures

All 14 visually impaired participants and 16 of the 250 sighted participants (mean age, 30.2 ± 5.1 years, range: 23–42 years) participated in the following functional magnetic resonance imaging (MRI) (fMRI) experiment. We measured brain activity using the same scanner when all blindfolded participants imagined that they were actually walking or jogging around a circle of 2 m radius, which was previously experienced on a different day (training). The training was conducted in a gymnasium, where blindfolded participants hearing white noise performed walking and jogging around a circle of 2 m radius (for more details, see Amemiya et al. (2021)). In this training, they were required to walk (or jog) along the circle as accurately as possible and return to the starting position. Therefore, this task requires spatial navigation without vision.

Before entering the MRI scanner, all participants (including the visually impaired participants) wore a pair of eye patches and an eye mask to eliminate any possible visual stimuli, which were maintained throughout the experiment. Their brains were scanned approximately 15–20 min after visual deprivation. This was consistent across participants, as eye closure duration greatly affects activity in the visual areas (Weisser et al., 2005; Merabet et al., 2007). Subsequently, the participants laid on the MRI scanner. Their heads were immobilized using sponge cushions, and their ears were plugged. Both arms were naturally semipronated and extended in front of the participants. Next, they were instructed to relax their entire body without producing unnecessary movements and to think only about things relevant to the tasks assigned.

Each participant completed two experimental runs. Each run comprised eight imagery epochs, each lasting 20 s. The imagery epochs were separated by 10-s baseline periods. Each run also included a 20-s period before the start of the first epoch. In one epoch, participants were asked to imagine either clockwise or counter-clockwise walking or jogging, resulting in four imagery conditions. Each condition was performed two times per experimental run in a pseudo-randomized order. During the experimental run, the participants were given auditory instructions (e.g., clockwise, walk, and start) through

MRI-compatible headphones to inform them of the action to be imagined and the starting time. We also provided the instruction “stop” to notify the participants of the cessation time for each epoch. The instructions were generated using a computer. Before the fMRI experiment, the participants were explicitly instructed to imagine the kinesthetic motor imagery of the experienced actions from a first-person perspective.

#### fMRI data acquisition

Functional images were acquired using T2*-weighted gradient-echo echo-planar imaging (EPI) sequences obtained via a 3.0-Tesla MRI machine (Trio Tim; SIEMENS, Germany) and a 32-channel array coil. Each volume consisted of 51 slices (slice thickness, 3 mm) acquired to cover the entire brain using multiband imaging (multiband factor 3)(Moeller et al., 2010). The imaging parameters were as follows: repetition time (TR), 1000 ms; echo time (TE), 27 ms; flip angle, 60°; field of view, $192 \times 192$ mm$^2$; matrix size, $64 \times 64$ pixels; and voxel size, $3 \times 3 \times 3$ mm$^3$. In total, 260 volumes were collected per run.

#### Imaging data analysis

fMRI data were preprocessed using the Statistical Parametric Mapping 8 (SPM8) toolbox (Department of Imaging Neuroscience, Institute of Neurology, UCL, London, UK), as previously reported (Amemiya et al., 2021). After preprocessing, we performed gPPI analysis using Conn toolbox version 21.a. The left RSC/PCC region-of-interest (ROI) (see the main text) was used as the seed region. For each participant, the time course of the average fMRI signal across the voxels in each ROI was deconvolved using the canonical hemodynamic response function (physiological variable). Next, we performed a general linear model analysis using the design matrix and included the following regressors: physiological variable, boxcar function for the task epoch (psychological variable), and multiplication of the physiological and psychological variables (PPI). These variables are convolved with a canonical hemodynamic response function. Six realignment parameters were included in the design matrix as regressors of no interest.

We generated an image of the voxels in which activity changed with the PPI regressor in each participant. These individual images were used in the second-level analysis. Here, we examined whether activity in the SPL, PHC, or visual cortices of a visually impaired participant enhanced its functional coupling with that of the left RSC/PCC ROI compared to the 16 sighted participants. We performed a two-sample non-parametric permutation test (without variance smoothing), which did not rely on the assumptions of a normal distribution and equal variance. This test was performed using the Statistical non-Parametric Mapping (SnPM) toolbox (version 13.1.09; Nichols and

Holmes, 2002). For statistical comparisons, we adopted 17 permutations based on a control group size of 16. Age and handedness scores were entered into the design matrix as covariates of no interest to minimize the impact of these variables on the findings.

In each visually impaired participant, we first generated a cluster image of voxels with a height threshold of $p < 0.005$ and evaluated whether a significant cluster was identified in the SPL, PHC, or visual cortices. To define the SPL and visual cortices, we used cytoarchitectonic probability maps of the Montreal Neurological Institute (MNI) standard brain in the SPM anatomy Toolbox v3.0 (Eickhoff et al., 2005; Amunts et al., 2020). The SPL ROI was defined as the bilateral cytoarchitectonic areas 5 (5Ci, 5L, and 5M) and 7 (7A, 7M, and 7PC). The visual ROI was defined as the bilateral cytoarchitectonic areas V1 and V2. For the PHC, the bilateral PHC ROI (see main text) was used. Each ROI image was used as an explicit mask to search for significant clusters of voxels in the SPL, PHC, and visual ROI (extent threshold of $p < 0.05$, FWE-corrected).

Results and Discussion

Among the 14 visually impaired participants, BS1, BNS5, and BNS7 showed a cluster with a significant trend ($P = 0.059$, corrected) within the SPL ROI. Similarly, these clusters were identified within the visual ROI in BS1 and BNS2. Such clusters were identified within the PHC ROI only in BS2. Therefore, only BS1 showed such clusters in the early visual cortices (peak coordinates x, y, and z = -8, -84, and 16 in area hOc1[V1]; T = 5.37; 65 voxels) and right SPL (peak coordinates x, y, and z = 14, -52, and 74 in area 5 L; T = 7.17; 115 voxels), in which activity enhanced functional coupling with that of the left RSC/PCC ROI during the imaginary spatial navigation task, compared to the sighted participants (Supplementary Fig. 3). The majority of voxels in the right SPL cluster (100 of the 115 voxels) were located in area 5. BS1 did not exhibit any significant clusters in the PHC.

It appears that when BS1 performs mental walking or jogging, the RSC/PCC likely communicates with the SPL (area 5) with an egocentric framework (Scheperjans et al., 2005; 2008; Naito et al., 2008), and the visual cortices innately have an allocentric (retinotopic) framework (Striem-Amit et al., 2015). This finding corroborates the view that the RSC/PCC is situated in an appropriate location for the association between egocentric and allocentric information.

**Voxel-based morphometry analysis for the hippocampus**

Based on previous findings (Fortin et al., 2008; Leporé et al., 2009), we also examined the increase in GM volume in the hippocampus in the visually impaired group (n = 14) and each visually impaired participant (n = 1) compared to the sighted group (n = 250). Using the Automatic Anatomical Labelling Atlas (AAL; Tzourio-Mazoyer et al., 2002), we defined the left and right 'hippocampus' regions of the AAL as the left and right hippocampal ROIs, respectively. We used the same nonparametric analysis and thresholds as in the main text.

As previously reported, the visually impaired group (n = 14) showed a significant GM volume increase within the right hippocampal ROI (peak coordinates x, y, and z = 15, -11, and -18; T = 3.34; 86 voxels), compared to the sighted group (Supplementary Fig. 4). This increase was mainly observed in the anterior part of the hippocampal ROI, which is consistent with previous findings (Fortin et al., 2008).

In the single-case voxel-based morphometry analysis, BS4 showed significant clusters within the left (peak coordinates x, y, and z = -20, -30, and -6; T = 4.35; 221 voxels) and right (peak coordinates x, y, and z = 26, -23, and -11; T = 4.25; 166 voxels) hippocampal ROIs. Similarly, BNS3 showed a significant cluster in the right hippocampal ROI (peak coordinates x, y, and z = 27, -39, and -2; T = 3.81; 241 voxels). However, BS1 did not show significant clusters within the left or the right hippocampal ROI. Eventually, the probability of significant clusters within the left and right ROIs (2/12) in the BS group was not significantly different from that (1/16) in the BNS group (Fisher's exact test, p = 0.56).

**Supplementary Figures and Figure Legends**

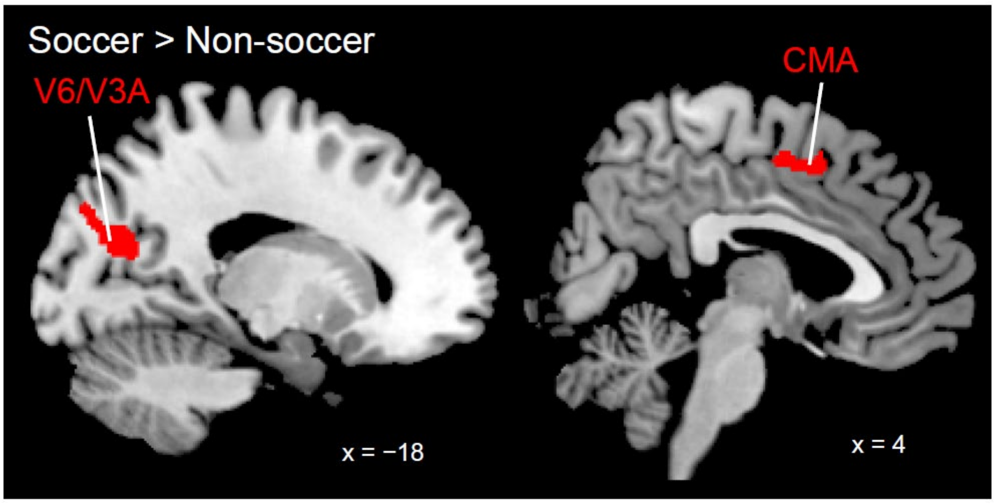


**Supplementary Fig. 1** Brain regions in which gray matter (GM) volume increased in the individuals with soccer experience for more than 4 years (n = 53) compared to those with no soccer experience (n = 160). These are superimposed on the MNI standard brain, and the sagittal slices (x = -18 and 4) are shown. MNI, Montreal Neurological Institute; CMA, cingulate motor area.

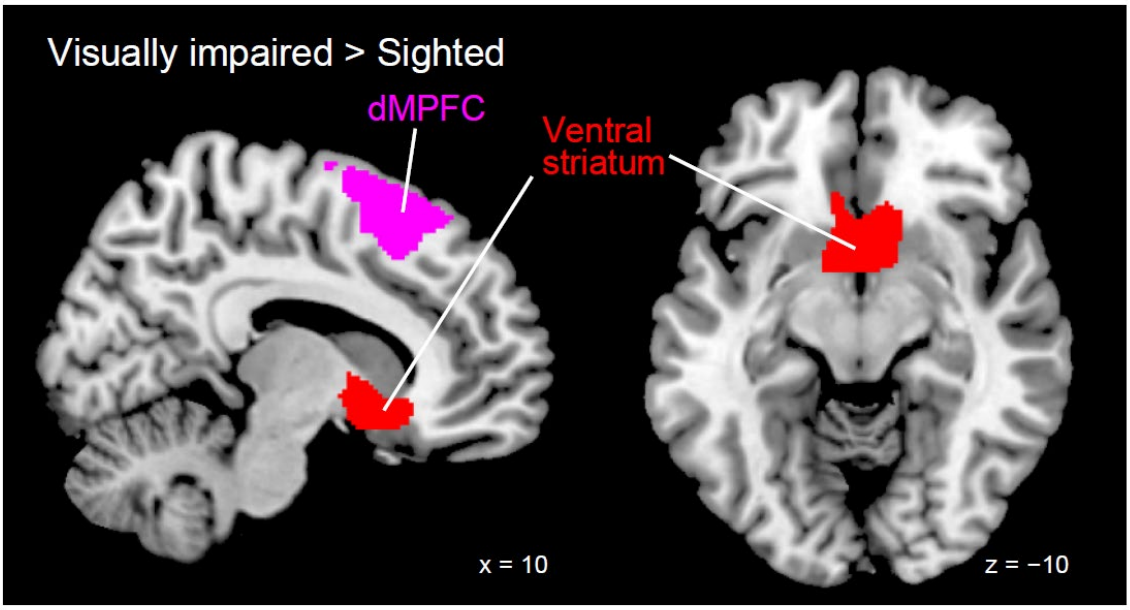


**Supplementary Fig. 2** Brain regions in which gray matter (GM) volume increased in the visually impaired group (n = 14) compared to the sighted group (n = 250). The ventral striatum cluster (shown in red) survived extent threshold with whole-brain correction ($p < 0.05$ FWE-corrected), while the dMPFC cluster (shown in pink) did not reach the threshold (but the second largest cluster). The regions are superimposed on the MNI standard anatomical image (left panel: sagittal slice of x = +10, right panel: transverse slice of z = -10. dMPFC, dorsomedial prefrontal cortex; FWE, family-wise error; MNI, Montreal Neurological Institute.

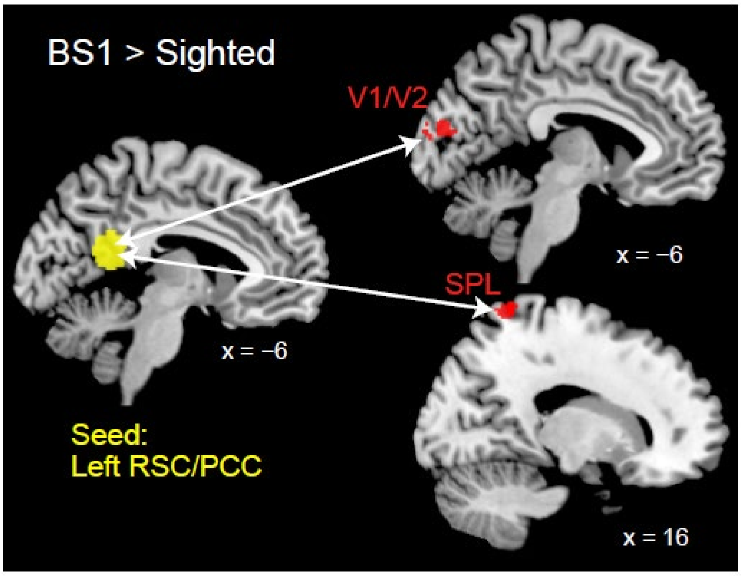


**Supplementary Fig. 3** Early visual cortices (V1/V2) and superior parietal lobule (SPL, area 5) (red sections) in which BS1 showed significantly higher functional connectivity with the left RSC/PCC ROI (yellow section) than the sighted participants during the imaginary spatial navigation task. These are superimposed on the MNI standard brain, and the sagittal slices (x = -6 and 16) are shown. BS, blind soccer; MNI, Montreal Neurological Institute; PCC, posterior cingulate cortex; ROI, region-of-interest; RSC, retrosplenial cortex.

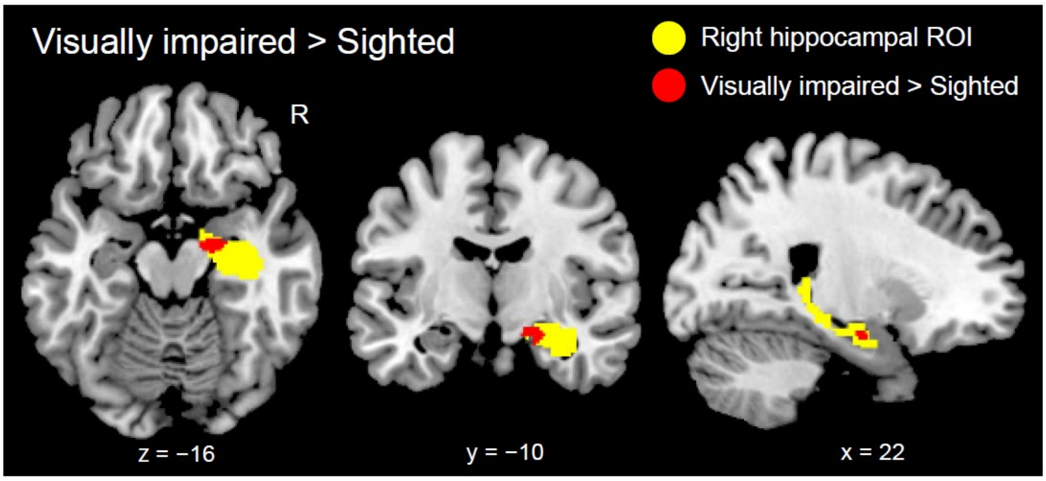


**Supplementary Fig. 4** GM increase within the right hippocampal ROI in the visually impaired group (n = 14) compared to the sighted group (n = 250). This is superimposed on the MNI standard brain, and the horizontal (z = -16), coronal (y = -10), and sagittal (x = 22) slices are shown. GM, gray matter; MNI, Montreal Neurological Institute; ROI, region-of-interest.

**Supplementary Table 1**

| ***RSC/PCC ROIs*** | *Significant cluster* | | |
|---|---|---|---|
| | Yes | No | Total number |
| *BS group* | 4 | 8 | 12 |
| *BNS group* | 0 | 16 | 16 |

Note: Significant group difference (Fisher's exact test, $p = 0.02$). BS, blind soccer; BNS, non-blind soccer; RSC, retrosplenial cortex; PCC, posterior cingulate cortex; ROI, region-of-interest.

**Supplementary Table 2**

| ***PHC ROIs*** | *Significant cluster* | | |
|---|---|---|---|
| | Yes | No | Total number |
| *BS group* | 1 | 11 | 12 |
| *BNS group* | 0 | 16 | 16 |

Note: No significant group differences (Fisher's exact test, $p = 0.43$). BS, blind soccer; BNS, non-blind soccer; PHC, parahippocampal cortex; ROI, region-of-interest.
